# Multimodal mismatch responses in associative but not primary visual cortex support hierarchical predictive coding in cortical networks

**DOI:** 10.1101/2023.04.12.536573

**Authors:** Alice B Van Derveer, Jordan M. Ross, Jordan P. Hamm

## Abstract

A key function of the mammalian neocortex is to process sensory data in the context of current and past stimuli. Primary sensory cortices, such as V1, respond weakly to stimuli that typical in their context but strongly to novel stimuli, an effect known as “deviance detection”. How deviance detection occurs in associative cortical regions that are downstream of V1 is not well-understood. Here we investigated parietal associative area (PTLp) responses to auditory, visual, and audio-visual mismatches with two-photon calcium imaging and local field potential recordings. We employed basic unisensory auditory and visual oddball paradigms as well as a novel multisensory oddball paradigm, involving typical parings (VaAc or VbAd) presented at p=.88 with rare “deviant” pairings (e.g. VaAd or VbAc) presented at p=.12. We found that PTLp displayed robust deviance detection responses to auditory-visual mismatches, both in individual neurons and in population theta and gamma-band oscillations. In contrast, V1 neurons displayed deviance detection only to visual deviants in a unisensory context, but not to auditory or auditory-visual mismatches. Taken together, these results accord with a predictive processing framework for cortical responses, wherein modality specific prediction errors (i.e. deviance detection responses) are computed in functionally specified cortical areas and feed-forward to update higher brain regions.

## Introduction

Context modulates information processing in the mammalian neocortex. A key effect of this contextual modulation is the suppression of responses to expected or contextually redundant stimuli ^1–7^ and amplification of responses to unexpected or contextually deviant stimuli ^8–12^. These phenomena have been studied using sensory “oddball” paradigms and termed stimulus-specific adaptation (SSA) and deviance detection (DD), respectively. Robust SSA and DD have been identified in visual, auditory, and tactile modalities across species^9, 11, 13^, suggesting that they represent universal computations in mammalian sensory cortex. In particular, DD has significant clinical relevance as well, as its EEG analogue “mismatch negativity” has been reliably shown to be reduced in diseases like schizophrenia^14^, pointing at a fundamental disruption in the brain circuits responsible for contextual processing^15^.

In primary sensory cortices, DD has been explained theoretically as a consequence of “predictive processing” in hierarchically organized cortical networks^10^, namely as a neural signature of a “prediction error”. Putatively under this framework, the brain generates internal predictive models, held mainly in higher brain areas, based on previous experience^16, 17^. These models are then compared against incoming sensory information in lower brain areas. This comparison is theoretically carried out via long-range feedback modulation from higher to lower areas^18, 19^, synapsing in layer 1. When internal models accurately predict bottom-up sensory inputs, responses in early sensory brain areas are suppressed, but when information deviates from said models, responses are amplified and sent upward in the hierarchical network as “prediction errors” to update internal models^18, 19^. This proposed process limits the amount of incoming sensory information the brain needs to process at each level of the cognitive hierarchy, thereby conserving time and energy^16^.

DD is well documented at the level of primary sensory regions to simple, unisensory stimuli. In primary visual cortex (V1), DD to visual oddball stimuli is present in layer 2/3 neurons, where bottom-up sensory data is integrated with top-down modulation from higher brain areas^11, 13^. This is consistent with DD representing a form of prediction error. Whether and how different forms of DD present across higher sensory processing brain areas is unknown but is nevertheless critical information in determining whether cortex is a predictive processing circuit. Past work shows that while low-level, unisensory DD is present lower cortical regions (e.g. primary auditory cortex; A1), it also propagates to higher brain regions^20–22^. This conforms with the predictive coding hypothesis, since prediction errors generated in early sensory regions should be fed-forward to hierarchically higher cortical areas to update internal models^16^. But it remains unclear whether a stimulus that deviates from contextual regularities only in its high-level properties – like a face or an audiovisual mismatch – elicits DD in low-level regions or only in higher-level, associative regions. If predictive processing theories generalize, then one may expect the latter, as prediction errors (DD) generated in a given cortical area should mainly propagate forward, not backward, in the heirarchy. In other words, higher associative regions may express DD to both low-level and high-level stimulus properties while lower-level areas, such as sensory cortices, may express DD solely to lower-level properties – i.e. the type of stimuli that region is functionally specialized to process, based on the bottom-up inputs it receives (arriving in layer 4, from the thalamus or lower cortical regions)^16, 18^.

If this is true, associative cortices in e.g. parietal regions should exhibit robust DD to unexpected multimodal stimuli, but lower regions like V1 should not exhibit DD to such stimuli if there is no deviance in the visual domain alone. To test this important postulate of the predictive coding hypothesis, we examined multisensory and unisensory DD in awake mice in V1 as well as parietal associative area (PTLp), a high-level visual area in mice^23^ often equated to posterior parietal cortex. PTLp is site of multisensory audio-visual integration, receiving inputs from both visual and auditory cortices and synthesizing them to interpret the environment in context ^24–27^. We employed a typical oddball paradigm (deviants =p.12) for the unisensory portion, and, for the multisensory portion, we designed a novel paradigm producing exclusively “multisensory” deviants. Visual stimuli (Va or Vb) were paired with auditory (Ac or Ad) such that mice were habituated to VaAc or VbAd, and then presented mismatched pairings (VaAd) at rare probabilities (p=.12; Figure 1). This design does not allow auditory or visual components of this multisensory oddball paradigm to produce DD on their own, as there were only 2 auditory and 2 visual stimuli presented, and each occurred at p≈0.5. Thus, the deviants were exclusively “multisensory”, distinguishing our approach from some past work using dual auditory-visual oddball stimulus sequences in which deviance was present in both auditory and visual domains independently^28^. This enabled us to address the question of how integrated multisensory information is processed in context. We show that mouse PTLp displays robust multisensory DD, as expected, as well as visual DD. In contrast, V1 did not exhibit multisensory DD, but did exhibit strong unisensory visual DD. Neither region exhibited auditory DD. This pattern of results supports a generalized predictive processing model of sensory cortical responses.

**Figure 1:**
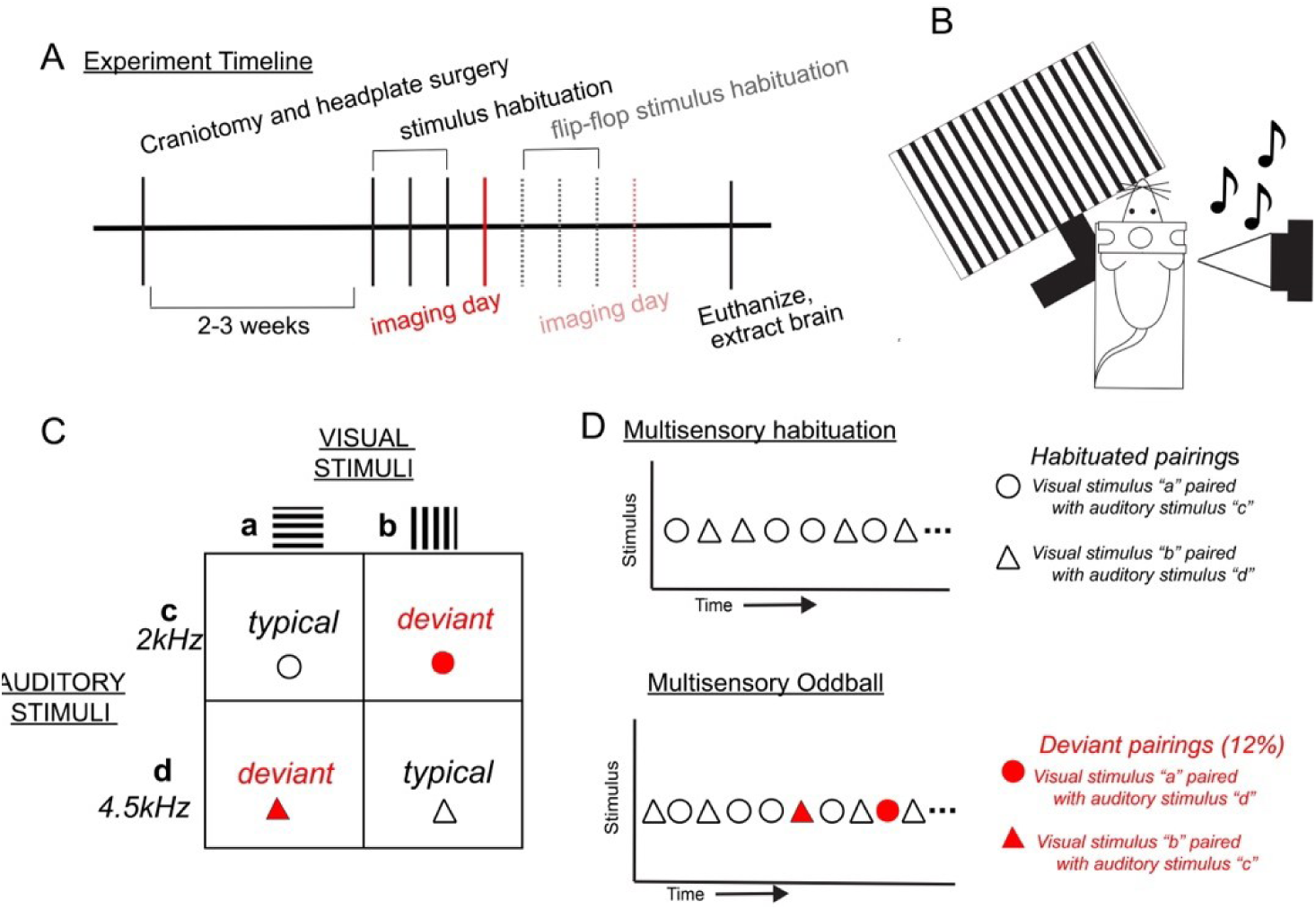
Paradigm for studying multisensory deviance detection in mice. A) Timeline detailing experimental events. B) Schematic of head fixation and stimulus presentations during habituation and imaging days. During these multisensory studies, mice were presented with (C) consistent auditory-visual pairs classified as “typical” (VaAc or VbAd) or “deviant” (VaAd or VbAc) for up to 30 minutes per day. (D) In the three days prior to the imaging day, mice were exposed to multisensory habituation runs consisting only of typical pairs. On imaging days mice were first exposed to two five-minute runs of the typical multisensory pairing and then presented with a “multi sensory oddball” run consisting of typical pairings (88% of trials) pseudorandomly interspersed with mismatched (i.e. “deviant”) pairings (12% of trials).

## Results

### Associative cortex detects deviant multimodal stimuli

We studied responses to multisensory deviants in awake mouse cortex. Animals were head-fixed and presented with audiovisual stimuli for three habituation days (Figure 1A), These stimuli consisted of two pairs, randomly presented in 2-3 runs of 6-7 minutes long: visual stimulus A (Va) with auditory stimulus C (Ac) or visual stimulus B (Vb) with auditory stimulus D (Ad). Visual stimuli were moving squarewave gratings (100% contrast, 8 cycles per degree, 2 cycles per second) at orthogonal orientations (0-180deg; randomized across mice) and auditory stimuli were 40-Hz amplitude modulated pure tones of either 2kHz or 4.5kHz. Stimuli were presented semi-simultaneously (<40ms offset) for 500ms in duration (Figure 1B,C). On the imaging days, animals were presented with a training stimulus run and a deviant stimulus run (Figure 1D). The majority of the stimuli (88%) in the deviant run were the same as during the habituation runs (VaAc; VbAd—the “typical” pairs), but occasionally, mismatched parings (Vb Ac; Va Ad) were presented (i.e the “deviant” pairs).

Although studies often include an active behavioral task during sensory processing paradigms as a strategy for ensuring attention to the stimuli^29^, we specifically excluded it and overt behavior, instead employing passive paradigms, for multiple reasons. First, animals are naturally able to detect unexpected stimuli in the absence of reward anticipation. Studying this function was an aim of our work. Second, recent work has shown that processing rewards or punishments activates top-down circuitry cortex-wide^30^. This could confound our results, as we aim to study region-specific responses to specific types of stimuli as they are processed in a bottom-up fashion.

To assess PTLp activity to multisensory deviants, we studied excitatory neuronal responses in transgenic mice expressing GCaMP6s, and imaged calcium activity in layer 2/3 (as layer 2/3 is known to exhibit the most robust DD responses in primary sensory regions ^11, 13, 31^). A cohort of VGlut/GCaMP6s transgenic mice (n=13; 11 female) experienced our multisensory oddball paradigm. We isolated 332 PYRs in PTLp showing strong responses to our multimodal stimuli (27.2% of PTLp neurons imaged; Figure 2A,B). Responses of PTLp neurons to deviant pairs were enhanced compared to typical pairs (Fig. 2c-e; (response to deviant vs typical pair, normalized for number of trials; paired t-test t(330)=3.85, p<.001).

**Figure 2:**
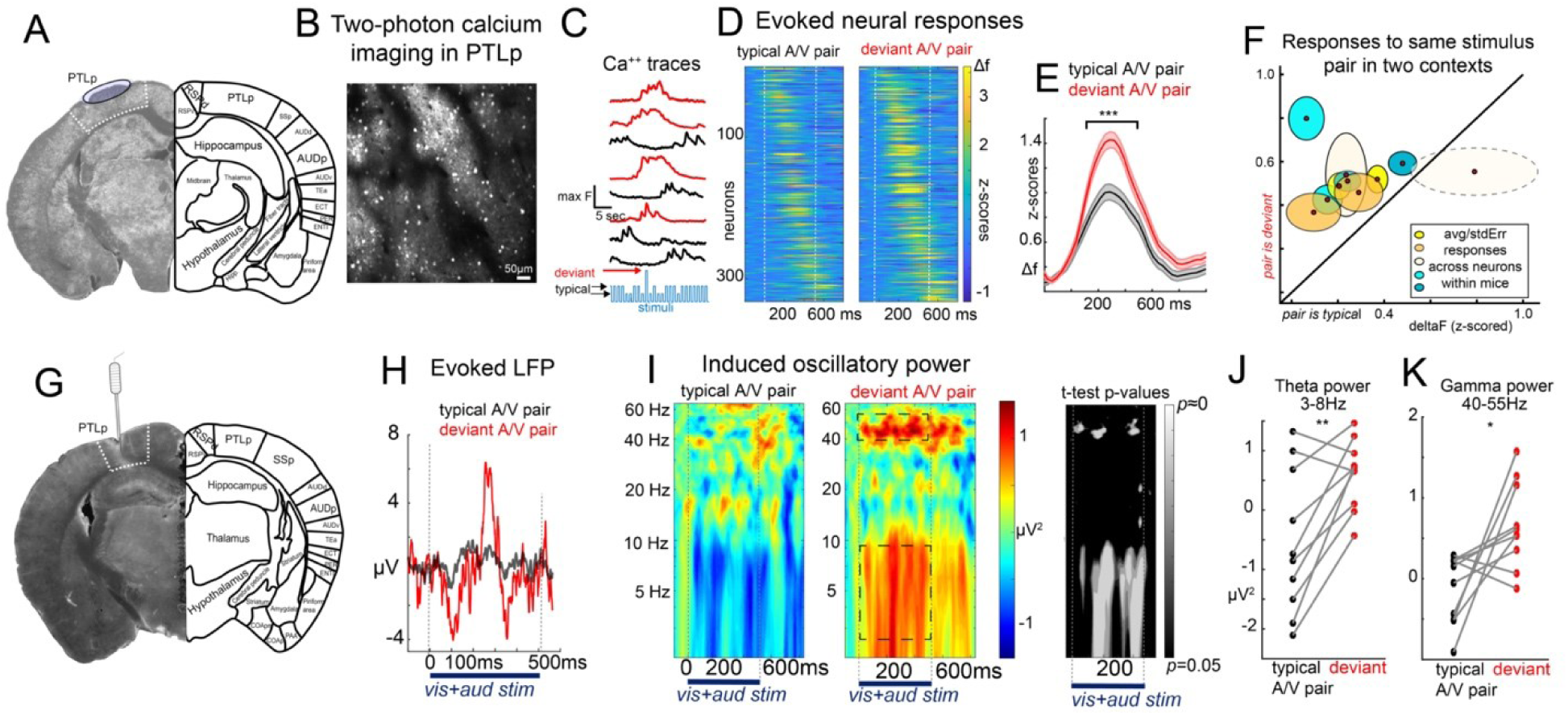
Associative cortex (PTLp) exhibits augmented responses to deviant multisensory pairs. A) Imaging took place in stereotaxically defined and confirmed PTLp according to the Allen Institute Mouse Brain Atlas. B) Example two-photon calcium imaging field of view in transgenic mice expressing GCaMP6s in L2/3 PTLp excitatory PYRs. C) Extracted calcium traces from isolated cells during the multisensory oddball paradigm D) reveals strong responses to the multisensory deviants relative to the previously habituated typical pairing. E) Paired samples t-test evinces significant “deviance detection” across neurons. F) A subset of mice (n=5) repeated the paradigm, re-habituating to the “deviant” A/V pair as the “typical” pair after the first imaging day (to rule out effects of feature selectivity). Within-mouse average responses (centroids) and within-mouse standard error (ovals) across responsive PTLp neurons evince strong deviance detection in all mice to both stimulus pairs except in one mouse, which only showed DD to one pair. G) Local field potential recordings (LFPs) in PTLp reveal H) a strong evoked response to deviants relative to all typical stimuli. I) Time-frequency analysis of single trial data shows a strong J) theta-band and K) gamma-band response during the stimulus period to deviant stimulus pairs relative to typical pairs (each point is one mouse). *p<.05,**p<.01,***p<.001.

The typical pairs and the deviant pairs consisted of different stimuli, and if PTLp shows highly specific multisensory selectivity, it could be possible that the differences we saw here were due to differences in feature selectivity, rather than genuine deviance detection. To rule this out, we carried out an additional “flipped” paradigm, re-habituating mice to the “deviant” pairs for three days following the first experimental recording, and then recording again on a 7^th^ day (Figure 1A, S2), so that PTLp responses could be quantified to the same stimulus pair when it was deviant vs typical. Neurons continued to display increased responses when the multisensory stimulus pairs were flipped (Figure 2F, Figure S2B). Only one mouse’s population responses to one of the paired showed no difference between the typical and deviant contexts (Figure 2F), suggesting that this was a robust phenomenon and confirming that the multisensory DD responses are not contingent on the pairings themselves, but rather the predictability of the paired stimuli.

DD is thought to reflect sensory prediction errors in the cortex which are fed-forward to higher brain regions. Past work has shown that feed-forward circuits in the cortex occupy mainly low-theta (3-8Hz) and gamma (>40Hz) frequency bands, while feed-back modulation occupies alpha and beta bands (10-30Hz)^32, 33^. If the DD we observed in PTLp represents a feed-forward signal (i.e. a prediction error) then it should present as increased theta- and gamma-band power specifically. In a separate cohort of mice, we analyzed local field potentials (LFP) recorded from PTLp during the same multisensory paradigm (Figure 1) to identify local circuit activity in response to predictable and unpredictable stimuli. We implanted bipolar electrodes into PTLp and recorded gross cortical responses during the multisensory oddball paradigm (n=10, females=5). We observed strong evoked responses to deviant relative to typical stimuli (Figure 2G,H). Additionally, time-frequency analysis of single trial data showed strong and statistically significant theta-band (Figure 2I,J) and gamma-band (Figure 2I,K) responses to deviant stimulus pairs compared to typical pairs (normalized for the number of trials; paired t tests theta: t(9)=3.73, p<.01; gamma: t(9)=2.82, p<.05). We did not observe main effects of sex or sex by context interactions for calcium imaging or LFP indices of DD (Figure S3).

Altogether, these results suggest that multisensory stimuli which deviate from expectations elicit DD signals in PTLp which may represent cortical prediction errors.

### Associative cortex displays deviance detection to unimodal visual stimuli

Our multisensory oddball paradigm was specifically designed to exclude unimodal deviance; that is, deviants were absent in purely the auditory or visual domain when taken in isolation (all auditory and visual stimuli were present at p=.5). So whether PTLp responds to unisensory deviants is not discernable from the above results. According to the theory of predictive coding, prediction errors should propagate upward in a hierarchically connected cortical network to update internal models of the environment. Although PTLp is a multisensory integration region, its anatomical inputs favor the visual stream^23, 34^, suggesting its fundamentally a higher order visual area which integrates auditory information (rather than an auditory area that integrates visual information). Thus, we hypothesized that PTLp should show unimodal deviance detection at least to visual stimuli.

To address this, a new cohort of VGlut/GCaMP6s transgenic mice (n=10; female=6) underwent two-photon calcium imaging in PTLp and were presented with standard visual or auditory oddball, flip-flop, and many standards control paradigms (serially presented, each ≈5 minutes) to examine single-neuron responses to repetitive and deviant unisensory stimuli in the form of full-field square-wave drifting gratings (visual; 0 or 90 deg drifting gratings) or pure tones (auditory; 2kHz or 4.5kHz). The unisensory paradigms enabled measuring responses when stimuli were rare and contextually deviant (oddball) or equally rare but not contextually deviant (many-standards control; Figure 3A,B), as previously described^35^. In PTLp, we isolated 192 PYRs (44% of total imaged) which responded to visual stimuli and 168 (38% of total imaged) which responded to auditory stimuli. As expected, we identified visual DD (control vs deviant: t(190)=2.29, p<.05), but not auditory DD (t(164)=-1.00, p=.32). Although visual DD was present in PTLp, it was a smaller effect size compared to the multisensory DD during the flip-flop paradigm (cohen’s D=0.33 vs 0.90).

**Figure 3.**
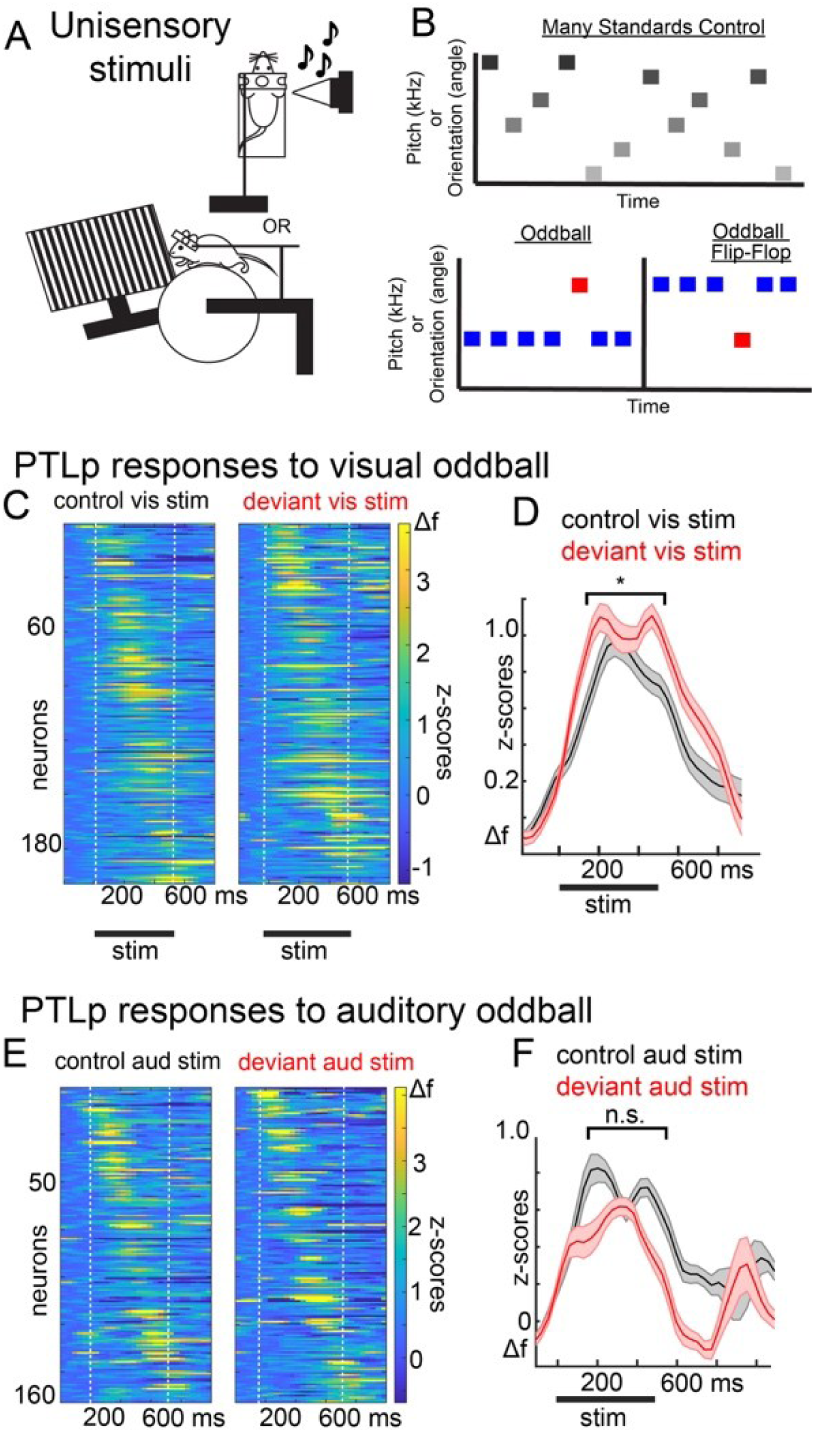
PTLp exhibits limited DD to unisensory deviants. A) Mice were presented with either visual or auditory stimuli in isolation in B) typical many standards and oddball paradigms. C) PTLp cortical neuron responses to control and deviant stimuli pooled across 10 mice (6 female). D) Averaged responses over 192 visually responsive PTLp neurons shows modest visual DD. E,F) same as C and D, but for auditory stimuli. Auditory stimuli evoked responses averaged over 168 PTLp neurons shows absent auditory DD. *p<.05

### Primary visual cortex detects visual deviance but not auditory or multisensory deviance

An additional extension of the hierarchical predictive processing hypothesis is that while primary visual cortex (V1) should detect deviance in line orientation – a feature for which V1 is strongly selective^36^ – it should not detect deviance for information that is fundamentally encoded or integrated downstream, such as audiovisual stimuli. That is, “oddball” stimuli that deviate from expectations only with regards to higher order properties should elicit prediction errors in higher brain areas (as seen above) but not necessarily lower regions.

We imaged layer 2/3 PYRs in V1 via two-photon calcium imaging during the same unisensory visual, unisensory auditory, and multisensory audiovisual stimuli in a separate cohort of mice. We isolated 109 layer 2/3 PYRs which responded to multisensory audiovisual stimuli (32% of V1 neurons recorded; n=4 mice, females=3) and 60 neurons which responded to unisensory stimuli (42% of V1 neurons recorded; as unisensory DD in V1 is a replication of various previous studies in this lab^11, 12, 35^, we opted to use a minimum number of mice (n=2; both female) for this experiment; Figure 4A,B).

**Figure 4:**
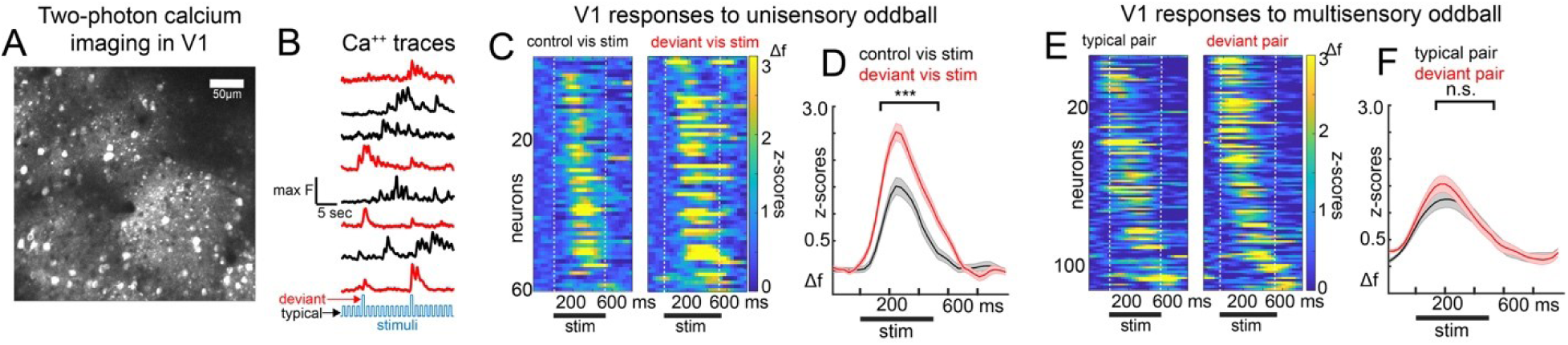
Primary visual cortex detects unisensory visual deviance but not multisensory deviance. A) Two-photon imaging in 6 mice in V1 viewing either the multisensory (4 mice) or unisensory (2 mice) oddball paradigms. B) V1 neuronal activity during a unisensory paradigm reveals C) augmented responses to deviant stimuli D) relative to many-standards control. E) V1 neuron activity to a multisensory paradigm revealed F) absent deviance detection, with typical and deviant pairs eliciting similar responses on average (109 responsive neurons, 32% of V1 neurons recorded). ***p<.01

As expected, audiovisual multisensory DD responses were not present in V1 (figure 4E,F; paired t-test multisensory: t(107)=0.71, p=.48), but, as previously shown, we observed strong unisensory DD to visual stimuli in V1 (Figure 4C,D; paired t-test unisensory: t(58)=3.10, p<.01). Additionally, while V1 neurons did show suprathreshold activity to auditory stimuli, evoked responses were small and did not exhibit DD (t(23)=0.03, p=.97). These findings suggest that, though projections both from auditory cortex to V1 and from PTLp to V1 exist, they do not convey DD responses to multisensory or auditory stimuli in V1. This is consistent with a hierarchical predictive coding framework, in which prediction errors are generated in layer 2/3 of regions of the cortex with selectivity for the deviant features, perhaps dependent on the integration of bottom-up inputs from the thalamus/layer 4 and top-down modulation arriving in layer 1.

## Discussion

Our results show that mouse PTLp, a higher-level associative region in the mouse visual cortical system, exhibits DD to contextually unexpected visual stimuli as well unexpected audiovisual stimuli. This multisensory DD was present in individual neuronal responses in layer 2/3, as well as increases in theta and gamma-band power in the LFP, consistent with the notion that this engaged feed-forward cortical circuitry^32^. In contrast, hierarchically lower V1 only exhibited DD to unexpected visual stimuli, but not to multisensory mismatches. Our findings are consistent with the hypothesis that predictive coding is an organizing principle of brain responses to the external world that generalizes across cortex^16^.

While EEG indices of visual DD such as P300 appear to propagate across the entire cortex^21, 37^, past data on whether individual neurons in higher brain areas beyond primary sensory cortices respond greater to deviant stimuli is inconsistent. We provide evidence that simple unisensory deviance propagates to higher brain regions and activates individual neurons. Interestingly, in our data, visual DD in PTLp was notably smaller than visual DD in V1 (almost 1/3^rd^ of the effect size). Due to differences in the paradigms (i.e. visual DD required a separate many-standards control run to isolate due to the impacts of stimulus specific adaptation^9, 35^, while multisensory DD in our paradigm was discernable in the same oddball run), we are cautious about direct comparisons in effect sizes between conditions. However, this is consistent with some past work on pure visual DD in ferret posterior parietal regions, which showed very modest visual DD in posterior parietal cortex^34^.

On the other hand, it is not necessarily consistent with findings in the auditory domain. Such work has shown stronger DD in prefrontal areas as compared to lower sensory cortices^20, 38^. As mentioned above, a pure interpretation of the predictive coding framework would suggest that locally generated prediction errors propagate to regions downstream (but not upstream). Still, V1 prediction errors may not flow from visual cortex exclusively to PTLp. For example, anterior cingulate area^39^ also receives strong V1 inputs, and is known to be involved in unisensory visual predictive processing^11, 12^. Thus, it is possible that there are different downstream circuits for different forms of visual predictive processing, and PTLp is less involved in pure visual orientation predictive modelling. Another possibility is that DD signals expressed in V1 are accommodated by areas hierarchically between V1 and PTLp, or by PTLp itself. Thus, the model updates before or at PTLp, and nothing is “fed forward” (hence, no DD or prediction error signal). Together, this could also explain why auditory DD was not identified in PTLp, despite it receiving inputs from auditory cortices. Above all, further work is needed to better understand the circuitry involved in the multisensory DD identified here, as well as how simpler, unisensory DD propagates across cortical hierarchies.

We did not identify multisensory DD in V1. This accords with a hierarchical view of predictive processing, in which an area should exhibit prediction errors (i.e. DD) only to stimuli for which it has feature selectivity. In canonical views of the predictive coding microcircuit, the convergence of top-down predictions arriving in layer 1 with bottom-up sensory data arriving from layer 4 or the thalamus should elicit prediction errors in subsets of layer 2/3 neurons^12, 18, 19^. Therefore, it is unlikely that V1 would have received the auditory information in the bottom-up direction, and, thus, it stands to reason that it could not generate canonical prediction errors to unisensory auditory or multisensory audiovisual deviants. Nevertheless, long range projections from auditory cortex do exist in V1, opening the possibility for multisensory integration to occur in V1 before being transmitted to areas of higher sensory processing, such as PTLp^40^. Recent work suggests, however, that auditory influence on V1 processing is modulatory and may arise simply from shared impacts of state or behavior^41^.

In apparent contrast with our findings, a previous study recording extracellular LFPs in anesthetized rats showed that purely auditory deviants do evoke an MMN-like signal in surface electrodes placed over V1, and visual deviants evoke an MMN-like signal over A1^28^, but this study did not measure responses of individual neurons. Further, auditory MMN in V1 took on the same temporal characteristics as auditory MMN in A1 (i.e. it was much earlier than visual MMN in V1, and vice versa). This could suggest that their findings reflect the synaptic inputs or modulatory input from one region to another, not the strong firing of layer 2/3 neurons to cross-modal deviants matching typical DD. Our finding of a lack of V1 DD responses to auditory deviants or to multisensory deviants conforms with this interpretation.

In conclusion, these results support theories of hierarchical predictive coding in the mammalian cortex and identify PTLp as a hub for audiovisual processing. Future work should determine whether the same pattern holds true for other combinations of sensory information in other associative regions. Further, whether the molecules, cells, and connections that support DD in V1 (i.e. interneuron functions, NMDAreceptors, layer 1 feedback inputs)^12, 19, 42^ are common to PTLp and other associative cortical regions should be investigated as well to depict how general the mechanisms of predictive processing are across the brain.

## Acknowledgments

This work was funded by the National Eye Institute (R01EY033950, Hamm), National Institute of Mental Health (K99/R00MH115082, Hamm; F32MH125445, Ross), Brain and Behavior Research Foundation (YI30149; Hamm), and the Whitehall foundation (2019-05-443; Hamm).

## Author Contributions

JPH and ABV designed the study, analyzed the data; All authors wrote and edited the manuscript; JPH supervised the work and acquired funding.

## Data availability statement

All data and code are available at https://gin.g-node.org/JordanHamm/Multisensory_paper.git or upon request to Jordan Hamm

## Materials and Methods

### Animals

All experimental procedures were carried out per the Georgia State University Institutional Animal Care and Use guidelines. Experiments were carried out in male and female transgenic mice expressing GCaMP6s in excitatory cortical neurons (two-photon calcium imaging) or wild-type C57BL/6 mice (local field potential). To create the transgenic animals, VGlut-cre animals (stock number 023527) were crossed with GCaMP6s reporter animals (stock number 031562) obtained from Jackson labs. Mice were maintained under a 12-hour light-dark cycle in a temperature and humidity-controlled environment with *ad libitum* access to food and water in the presence of environmental enrichment (tunnels, nesting material, etc.).

### Surgeries

Between P60 and P150, animals underwent surgery for head-plate fixation, followed by either the creation of a chronic cranial window (VGlut/GCaMP6s) or the insertion of bipolar electrodes (wild-type) (Fig. 1a). For head plate fixation, both types of mice were anesthetized with isoflurane (induction at 3%, maintenance at 1-2%). A titanium head plate with a circular opening in the center was attached to the skull with dental cement. For mice used in two-photon calcium imaging experiments, a small circle (3 mm in diameter) of skull was removed centered over left multisensory associative area PTLp (i.e. putative mouse parietal associative area, centered at -2.0 A/P and -1.7 M/L from bregma) or over left primary visual cortex (X =2 mm, Y = -2.92 mm) of VGlut/GCaMP6s mice and replaced with a 3-mm glass coverslip. The coverslip was held in place using dental cement. For the PTLp LFP experiments, a small hole was drilled at the PTLp coordinates mentioned above in wild-type mice. Bipolar titanium electrodes (Platstics One, Roanoke, VA, USA) were manually twisted together so that their contacts were separated by < 500µm and were gradually lowered to a depth of approximately 200 µm below the surface of the cortex. A common ground electrode was affixed to the skull with dental cement. Mice were allowed to recover in their home cage and given three days of analgesics (5 mg/kg carprofen intraperitoneally). Animals were accustomed to head fixation using three days of treadmill training, in which they were placed on a treadmill and head-fixed for 30 minutes at a time, but no data was collected. During training sessions and before the first imaging session, mice were exposed to either unisensory control stimuli (visual and auditory, separated) or multisensory training stimuli (combined visual and auditory). Locations of imaging and LFP recordings were confirmed via post-hoc histological verification via comparison with the Allen Institute Mouse Brain Atlas (Figure 2A,G) as previously described^11, 12, 35^.

### Unisensory Visual Stimulation

Visual stimuli were generated using the MATLAB (MathWorks) Psychophysics Toolbox and displayed on a liquid crystal display monitor (19-inch diameter, 60 Hz refresh rate) positioned 15 cm from the right eye, roughly at 45° to the long axis of the animal. They were presented with drifting grating stimuli consisting of a full-field square-wave grating (100% contrast, 0.04 cycles per degree, two cycles per second) drifting in eight different directions (0°, 30°, 45°, 60°, 90°, 120°, 135°, 150°) in random order. Stimuli were presented for 0.5 seconds with an interstimulus interval (ISI) of 0.5-0.6s. Mice underwent three training days with the random order of drifting gratings before data collection. Sequences of gratings were synchronized with two-photon imaging or electrical potential acquisition using PrairieView software (Bruker Inc, Billerica, MA, USA). For the visual oddball paradigm, two gratings that were perpendicular to each other were selected (0° and 90°, 45° and 135°, 30° and 120°). One grating served as the redundant stimulus and was presented 87.5% of the time, while the other served as the deviant stimulus and was presented 12.5% of the time. A flip-flop sequence was then shown in which the previous redundant stimulus became the deviant stimulus, and the previous deviant became the redundant. Each sequence had 250 trials with stimulus presentation for 0.5 seconds and a 0.5-0.6s ISI.

### Unisensory Auditory Stimulation

Auditory stimuli were generated using MATLAB (MathWorks, Natick, MA, USA) and presented through a multi-field magnetic speaker (Tucker-Davis Technologies, Alachua, FL, USA) within 3 cm of the mouse’s right ear. Mice were presented with pure auditory tones (0.5s stimulus duration, 0.5-0.6s ISI, 63dB). Eight different frequencies were presented randomly for the ‘many standards’ control (2.0 kHz, 3.0 kHz, 4.5 kHz, 6.8 kHz, 10.1 kHz, 15.1 kHz, 22.8 kHz, 34.2 kHz) sinusoidally amplitude modulated at 40Hz (100%). Mice underwent three training days with the many standards control before data collection. Sequences of tones were synchronized with image acquisition using PrairieView software. For the auditory oddball paradigm, one tone (2.0 or 4.5 kHz) served as the redundant and was presented 87.5% of the time, while a different tone served as the deviant stimulus and was presented 12.5% of the time. A flip-flop sequence was then presented in which the previous redundant stimulus became the deviant stimulus, and the previous deviant became the redundant. Each of these sequences had 250 trials with a presentation of 0.5s and an ISI of 0.5-0.6s.

### Multisensory Stimulation

The same timing parameters were employed for the multisensory condition (Figure 1B). Mice were presented with two combinations of an auditory (A) tone semi-simultaneously with a visual (V) drifting grating stimulus for 0.5s (ISI 0.5-0.6). The visual component consisted of a full-field square-wave grating (100% contrast, 0.04 cycles per degree) of either 45° (stimulus “Va”) or 135° (Vb). The auditory component consisted of either 2.0 kHz (Ac) or 4.5 kHz tones (Ad; Fig 1c). Sequences of auditory-visual stimulus pairings were synchronized with the image or electrical potential acquisition using PrairieView software. For the training stimulation, the same auditory tones were always presented with the same visual stimuli so that Ac was always paired with Va, and Ad was always paired with Vb (Fig 1d). Mice underwent three days of training sessions before the experimental session. For the experimental sessions, the sessions started with one run of the training pairs. Then, an “oddball” sequence was presented, involving a combination of auditory-visual pairings deviated so that deviants pairings were presented 12.5% of trials. That is, Aa was presented with Vb in 6.25% of the trials, and Ad was presented with Va in 6.25% of the trials (Figure 1D). Each of these sequences had 350 trials.

To ensure that we were measuring multisensory deviance detection and not feature selectivity to specific audio-visual combinations, one cohort of animals (n=7) was exposed to the above paradigm and a “flipped” multisensory paradigm (Figure S2a). Mice underwent three additional days of training sessions in which Va was always paired with Ad and Vb was always paired with Ac (the previously deviant combinations). The experimental session started with one run of the flipped training pairs and then the presentation of an “oddball” flipped sequence in which Va was paired with Ac in 12.5% of the trials, and Vb was paired with Ad in 12.5% of the trials. The responses to the original multisensory paradigm and the “flipped” multisensory oddball paradigm were compared.

### Two-Photon Calcium Imaging

On the day of imaging, awake mice were head-fixed on the treadmill to allow the mice to locomote freely throughout the experiment. The activity of cortical pyramidal cells in putative layer 2/3 (approximately 150-300 µm below the pial surface) was recorded by imaging fluorescence changes with a two-photon microscope (Bruker Ultima In Vivo; Billerica, MA) excited with a Ti:Sapphire laser (Chameleon Ultra II, Coherent) at 940 nm through a water immersion objective. The objective lens was immersed in ultrasound gel (Aquasonic Clear). Scanning and image acquisition were controlled by PrairieView software (Prairie Technologies; 28 frames per second, 256 x 256 pixels, 803 μm FOV)

### Calcium Imaging Analysis

Calcium imaging data sets were scored similarly to previous reports ^35, 43–45^. Raw images were processed to correct motion artifacts using the “Moco” plugin for ImageJ^46, 47^. Regions of interest (ROIs) were selected semi-automatically from time-averaged images using an in-house written MATLAB (MathWorks) routine^43^, and fluorescence traces were subtracted from the pixels just outside these ROIs (i.e., halo subtraction). Delta-f, i.e., the positive first derivative of LOWESS-smoothed fluorescent data (1 second window), normalized by the standard deviation of the lowest 8% of non-zero values, was calculated as a proxy for neuronal activity as previously described^43^. Delta-f was averaged across trials for each stimulus type (visual control, redundant, deviant; auditory control, redundant, deviant) and orientation (visual)/tone (auditory). We focused analyses on cells with stimulus-evoked responses >+2 standard deviations above pre-stimulus baseline on at least one stimulus type for at least one context/condition (i.e., they must be “stimulus-driven”). Trials containing significant motion artifacts were discarded from the analysis. Paired samples t-tests were carried out with individual cells as observations to test for DD by comparing the area under the curve between responses to control and deviant stimuli (two-tailed significance) for the unisensory paradigm, and between repsones to the typical pairs and the deviant pairs for the multisensory paradigm.

For the cross-week comparisons (figure 2F, S2) examining responses in the same mice/regions to the same stimulus pairs across weeks when they were typical vs deviant, we used independent t-tests, as we were not always able to identify exactly the same neurons. We also examined mousewise responses in the multisensory paradigm by averaging across all responsive neurons within each mouse to the typical vs deviant pairs in figures 2F and S2.

### Local field potential signal processing and analysis

Local field potentials (LFPs) were amplified with a differential amplifier (Warner instruments, DP-304A, high-pass: 0 Hz, low-pass: 500 Hz, gain: 1K, Holliston, MA, USA). Amplified signals were passed through a 60 Hz noise cancellation machine (Digitimer, D400, Mains Noise Eliminator, Letchworth Garden City, UK), which, instead of filtering, creates an adaptive subtraction of repeating signals which avoids phase delays or other forms of waveform distortion. Trials with excessive signal (>≈5 std devs) were manually excluded (between 0 and 20 per recording). LFPs were only collected in the multisensory paradigm. We used the third control stimulus after each deviant for generating comparisons (so that the number of trials were kept the same for both conditions^11^). This did not significantly impact our conclusions (Figure S1). Analyses were combined across both stimulus pairs. Ongoing data were converted to the time-frequency domain with a modified morelet wavelet approach with 100 evenly spaced wavelets from 2 to 70Hz, linearly increasing in length from 1 to 20 cycles per wavelet, applied every 10ms from 300ms pre to 700ms post stimulus onset (200ms post-stimulus offset) as previously described^35^. Stimulus induced power spectra were computed for both conditions for each mouse and baseline corrected by subtracting the average for each frequency in the 100ms prior to stimulus onset. Two peaks in the stimulus induced time-frequency power spectra were identified: one from 3 to 9Hz (theta) and one from 40-55Hz (gamma). We carried out paired t-tests on average power during the stimulus period for each mouse for each stimulus pair type (typical vs deviant) for each of these frequency bands.

