## Supplemental Figures S1-S3 for "Multimodal mismatch responses in associative but not primary visual cortex support hierarchical predictive coding in cortical networks"

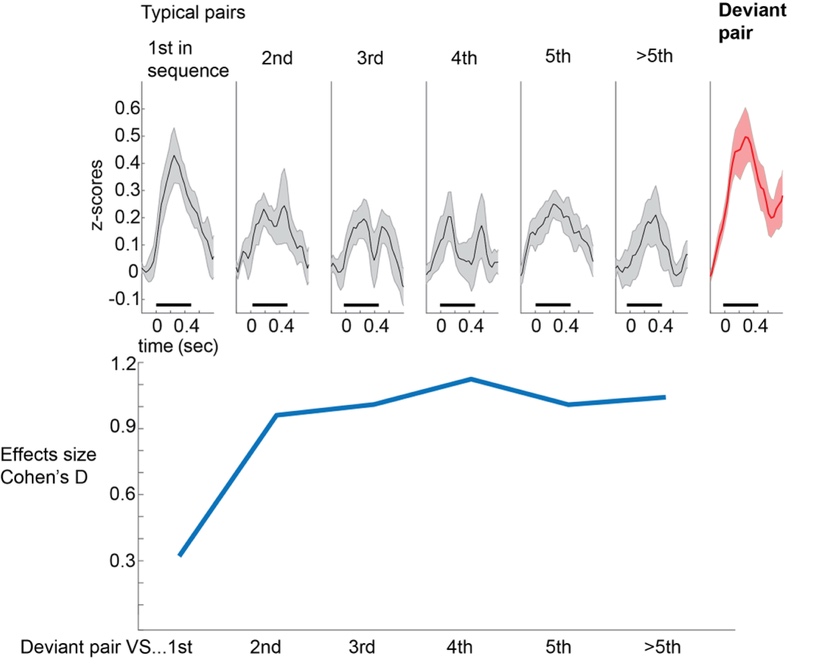

**Figure S1: Mousewise comparisons across typical stimuli in sequence.** Top: Responses across all responsive neurons (i.e. avg response >1std over baseline) were averaged all 13 mice to the typical stimulus pairs (according to their position in sequence after the first deviant) and to the deviant stimulus pair. Bottom: comparison of effect size (cohen’s D) comparing the deviant pair to each of the responses above. Some dishabituation was present after the presentation of the deviant, but this quickly resolved by the second stimulus, with effect sizes hovering around D=1.0. All comparisons in the main text used the 3^rd^ typical stimulus in the sequence.

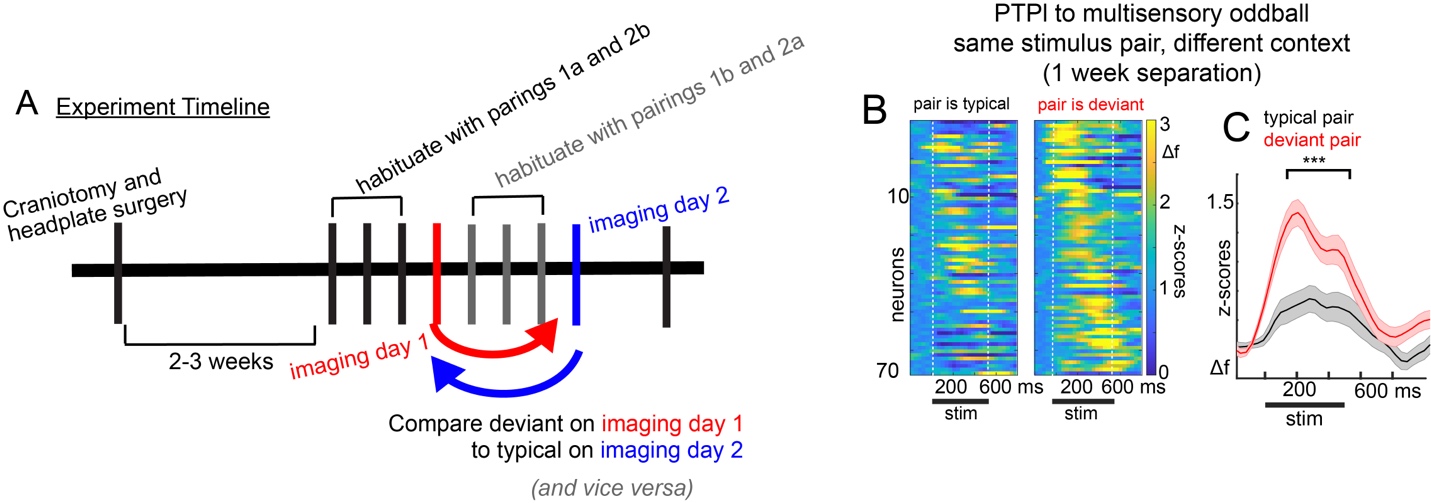

**Figure S2. Flip-flop paradigm – cell-wise comparisons.** A) A subset of mice habituated first to two auditory/visual stimulus pairs and then viewed an oddball paradigm on the imaging day with the pairings swapped on 12.5% of trials. The swapped pairings were habituated over three additional days, and the original pairings were presented as the “deviants” in the oddball sequence on imaging day 2. B) Comparisons between the original pairings when they were “typical” vs when they were “deviants” showed C) strong deviance detection to deviant pairs in PTLp (t(70)=3.76, p<.001), suggesting that simple stimulus pair or feature selectivity could not account for the main effects.

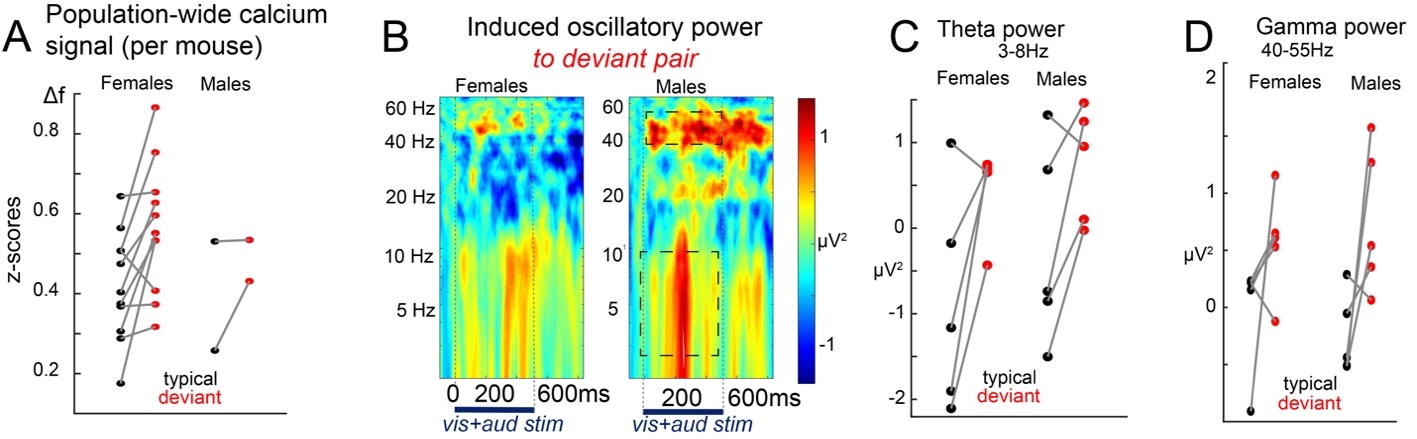

**Figure S3. Absent sex-differences in multisensory deviance detection.** A) Scatter-plot comparisons of average neural responses to typical vs deviant pairs during two-photon calcium imaging across all 13 mice suggest that males do not strongly differ from females in multisensory DD, although the number of males completing the two-photon calcium imaging study was not sufficient to statistically confirm this trend. Sufficient spread was collected in the local field potential experiments (n=10, 5 females). B) Induced oscillatory power to the multisensory deviants showed theta and gamma peaks in both males and females. Average spectra show ostensibly stronger responses in males overall, but C) theta and D) gamma power did not show sex main-effects (theta F(1,8)=1.00, p=.35; gamma F(1,8)=1.84, p=.21) or sex by stimulus context interactions (theta F(1,8)=2.21, p=.18; gamma F(1,8)=3.25, p=.11).
